# Short-term plasticity at retinogeniculate synapses depends on synaptic strength

**DOI:** 10.64898/2026.03.26.714222

**Authors:** Florian Hetsch, Irene Santini, Christina Buetfering, Sonia Ruggieri, Eric Jacobi, Jakob von Engelhardt

## Abstract

Relay neurons of the dorsal lateral geniculate nucleus (dLGN) receive convergent inputs from retinal ganglion cells (RGCs). Retinogeniculate synapses exhibit a highly skewed distribution of synaptic strength, with a few strong inputs and many weak ones. Strong synapses are thought to dominate relay neuron activity. However, the contribution of individual inputs might not just depend on strength but also on short-term plasticity Using minimal stimulation recordings in acute mouse brain slices, we analyzed the electrophysiological properties of individual retinogeniculate synapses. We observed a robust inverse correlation between synaptic strength and short-term plasticity: weak synapses showed facilitation, whereas strong synapses exhibited pronounced depression. This was consistent with increasing vesicle release probability and enhanced AMPA receptor desensitization at stronger synapses. Analysis of synaptic current kinetics further suggested that variability in synaptic strength reflects not only differences in synapse size and AMPA receptor content but also differences in the electrotonic distance of synapses from the soma. Together, these results reveal systematic heterogeneity in both presynaptic and postsynaptic properties of retinogeniculate synapses. Therefore, the relative contribution of weak and strong inputs to relay neuron firing is likely activity-dependent, with strong synapses dominating when RGCs fire few action potentials and weaker inputs contributing more during sustained or high-frequency firing with several action potentials.

## Introduction

Relay neurons of the dorsal lateral geniculate nucleus (dLGN) receive visual information from RGCs^1,2^. This information is not just relayed to the visual cortex, but is transformed by the active and passive properties of relay neurons and, in particular, by the dynamics of retinogeniculate synapses^3,4^. Classical functional studies in the cat and mouse visual system showed that dLGN relay neurons receive inputs from a few (1 to 3) very strong retinogeniculate synapses, often referred to as dominant inputs^5,6^. The strength of these synapses, with AMPA receptor-mediated current amplitudes ranging from hundreds of pA to several nA, is sufficient to cause relay neurons to fire^7-10^. However, *in vivo* analyses from cats with dual recordings from the retina and dLGN suggested that most of the relay neuron spikes cannot be attributed to single dominant inputs, but rather arise from the summation of excitatory postsynaptic potentials (EPSPs) from several weaker inputs^11,12^. Indeed, functional studies support the existence of weak connections, indicating that relay neurons can receive weak functional inputs from numerous RGCs^13,14^. Consistent with this, anatomical studies revealed that convergence might be even greater in mice, with up to 91 RGCs impinging onto a single relay neuron^15-17^.

The functional properties of the dominant inputs have been analyzed in great detail in numerous studies^9,13,18,19^. They form very large retinogeniculate synapses with multiple (approximately 30) release sites. A special feature of these synapses is their pronounced short-term depression during repetitive stimulation, which can be explained by a high vesicle release probability of 0,7^20,21^. Short-term depression in this synapse is further augmented by desensitization of postsynaptic AMPA receptors. The special geometry of these large synapses prolongs the glutamate dwell time in the synaptic cleft, promoting receptor desensitization. Importantly, spillover of glutamate from active to inactive neighboring release sites enhances the desensitization of AMPA receptors^21^. A slow recovery of AMPA receptors from desensitization is another factor responsible for short-term depression^21^. This can be explained by the expression of GluA1-containing AMPA receptors, which have a comparably slow recovery from desensitization and by the interaction of AMPA receptors with the auxiliary subunit CKAMP44, which effectively slows the recovery from desensitization^21-23^.

The numerous weaker retinogeniculate synapses have received much less attention than the dominant inputs. Almost nothing is known about their physiological properties.

Here, we characterized the electrophysiological properties of retinogeniculate synapses with a particular emphasis on the weaker synapses. We performed minimal stimulation of the optic tract to analyze the short-term plasticity of AMPA and NMDA receptor-mediated currents in acute brain slices from wildtype mice. We found an inverse correlation between synapse strength and short-term plasticity, with weak synapses facilitating and strong synapses depressing. Compound EPSCs elicited by maximal stimulation of the optic tract with a train of stimuli show initial depression followed by facilitation, suggesting that the contribution of strong and weak synapses changes dynamically during prolonged activation of retinogeniculate synapses. We used CKAMP44^-/-^ mice, in which AMPARs recover faster from desensitization, to investigate the contribution of desensitization to the inverse correlation between synapse strength and short-term plasticity. Our data suggest that presynaptic mechanisms (release probability) and postsynaptic mechanisms (desensitization) contribute to differences in short-term plasticity between weak and strong retinogeniculate synapses. Together, our results provide new insight into how diverse retinogeniculate synapses might shape thalamic processing and raise important questions about the functional role of weak retinal inputs in visual information transfer.

## Results

### Minimal stimulation reveals heterogeneity of retinogeniculate inputs strength

First, we characterized the properties of retinogeniculate synapses by performing whole-cell patch-clamp recordings from thalamic relay neurons in the dLGN in acute brain slices prepared from wildtype mice (P27-P50). To activate individual RGC inputs, we used minimal stimulation of the optic tract using a stimulation pipette with a monopolar electrode (Fig. 1A). Single fibre inputs had a median AMPAR-mediated EPSC amplitude of 47.84 pA (± 73.05 pA, median ± IQR, n=175). Notably, the distribution was strongly skewed toward small EPSC amplitudes (Fig. 1B, left). This is consistent with the previous observation that weak synapses outnumber strong inputs^13^. A similarly skewed distribution was observed for NMDAR-mediated EPSC amplitudes, with a median amplitude of 23.35 pA (± 43.16 pA, median ± IQR, n=103) (Fig. 1B, right). There was no correlation between AMPA/NMDA ratio and EPSC amplitude (Spearman’s correlation, r=0.052, p=0.601) (Fig. 1C), indicating that AMPA and NMDA receptor content scales proportionally with synaptic strength. The heterogeneity of EPSC amplitudes might result from the different distance of the synapse from the recording site, i.e., the cell body. If that were the case, smaller EPSCs would be expected to exhibit slower kinetics (rise and decay time), as these are also affected by dendritic filtering^24^. Indeed, we observed a negative correlation between the decay time constant τ_w_) and the amplitude of AMPAR-mediated EPSCs (Spearman’s correlation, r=-0.398, ****p<0.0001) (Fig. 1D). However, there was only a weak correlation between the 10–90% rise time and amplitude of EPSCs (Spearman’s correlation, r=0.244, *p=0.026) (Fig.1E), indicating that synapse location only explains a minor fraction of the observed variability in EPSC amplitude. The observed heterogeneity in synaptic strength and receptor kinetics is likely driven in part by differences in postsynaptic AMPAR content. Nevertheless, presynaptic factors, such as variations in vesicle release probabilities, may also contribute to the observed variability in synaptic strength.

**Figure 1.**
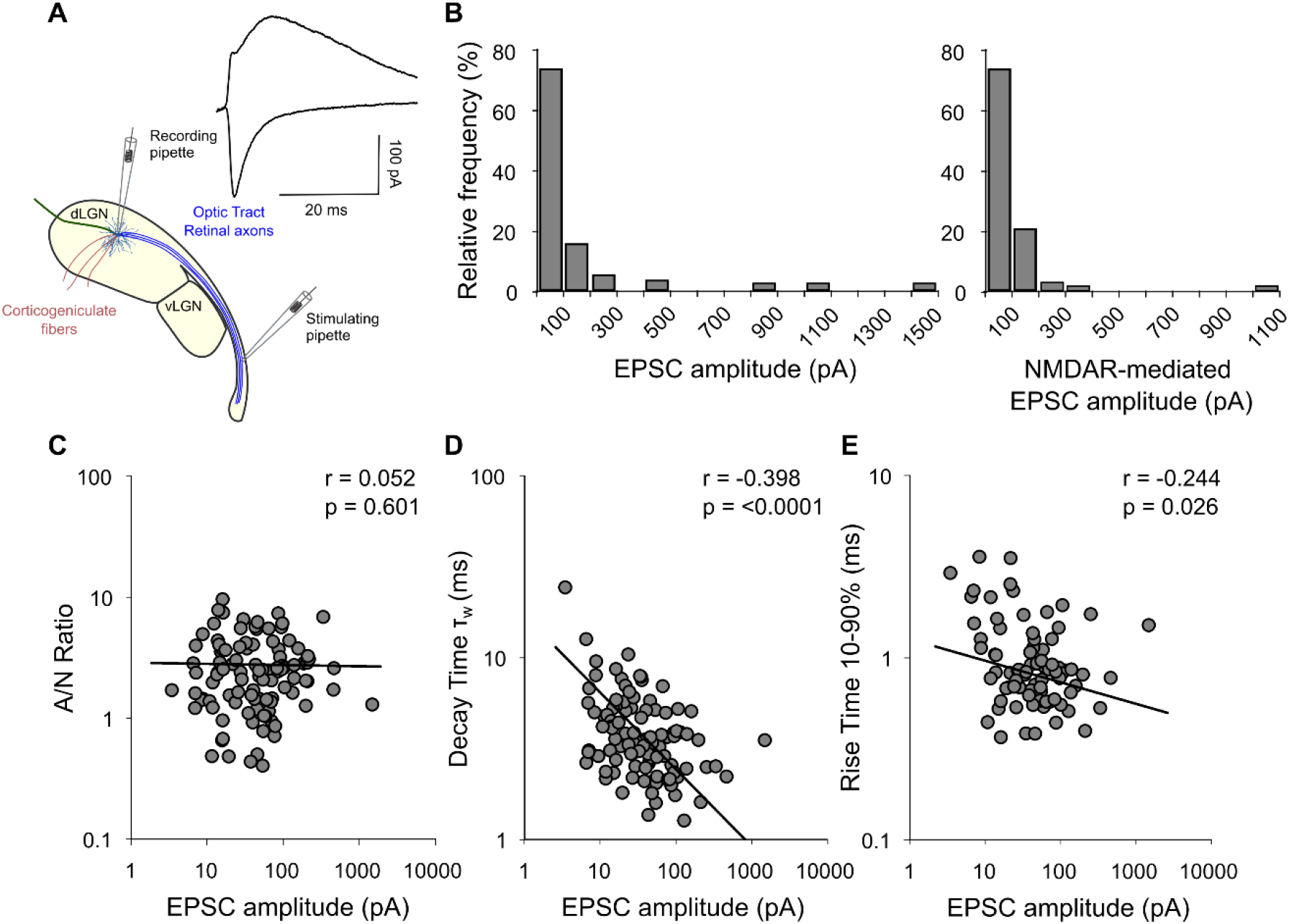
Characterization of the retinogeniculate synapse. (A) Schematic representation of the patch-clamp experiments and example currents. Whole-cell recordings were made from relay neurons in the dLGN while stimulating retinal axons in the optic tract with a monopolar electrode (minimal stimulation). (B) Distribution of the EPSC amplitudes across recorded inputs, ranging from tens of pA to >1 nA. AMPAR-mediated currents on the right (n=175); NMDAR-mediated currents on the left (n=103) (C) Dependence of AMPA/NMDA ratio on EPSC amplitude plotted on logarithmic scales (log Y vs. log X). (Spearman’s correlation, r=0.052, p=0.601, R^2^=0.0004, n=103). (D) Dependence of decay time constant (τ_w_) on EPSC amplitude plotted on logarithmic scales. Strong inverse correlation between τ_w_ and EPSC amplitude displays faster decay kinetics at stronger inputs and vice versa (Spearman’s correlation, r=-0.398, ****p<0.0001, R^2^=0.250, n=104). (E) Same as in (D), but for the rise time. Weak inverse correlation between 10–90% rise time and EPSC amplitude (Spearman’s correlation, r=-0.244, *p=0.026, R^2^=0.176, n=83). The data (black dots) are fitted by a linear regression (black solid line) in log–log space. The black line shows the best-fitting power law for the entire sample in each scatter plot.

### Inverse correlation between synaptic strength and short-term plasticity at retinogeniculate synapses

To investigate presynaptic function, we examined the short-term plasticity of retinogeniculate synapses. Driver inputs to thalamic relay neurons exhibit pronounced short-term depression^4,10,12^. However, less is known about the additional, weaker retinogeniculate synapses. To determine whether short-term plasticity correlates with synaptic strength, we performed minimal stimulation of the optic tract and recorded responses to trains of ten stimuli delivered at 50 Hz. We then quantified the PPR PPR_2/1_ (2^nd^ EPSC/1^st^ EPSC) and PPR_10/1_ (10^th^ EPSC/1^st^ EPSC) (Fig. 2A). We observed a strong inverse correlation between EPSC amplitude and PPR_2/1_ (Spearman’s correlation, r=-0.569, ****p=0.0001) or PPR_10/1_ (Spearman’s correlation, r=-0.479, **p=0.0015). Thus, consistent with previous observations^4,10,12^, strong “driver-like” synapses display short-term depression. In contrast, weak synapses facilitate (Fig. 2B). Since weak synapses outnumber strong synapses, retinogeniculate synapses showed, on average, short-term facilitation (Fig. 2C). Importantly, this does not imply that the net input from RGCs to relay cells facilitates. Because depressing inputs generate much larger EPSCs than facilitating synapses, compound responses composed of weak and strong inputs may display a different short-term plasticity than the average single fibre input recorded with the minimal stimulation approach. Indeed, when we activated multiple inputs simultaneously using maximal stimulation of the optic tract, compound EPSCs exhibited an initial paired-pulse depression (PPR_2/1_: 0.85 ± 0.31; mean ± SD, n=19), followed by weak facilitation at the end of the 10-pulse train (PPR_10/1_: 1.25 ± 0.306; mean ± SD, n=19) (Fig. 2D). This transition from depression to facilitation can be explained by the interplay of depressing strong inputs and facilitating weak inputs: strong inputs dominate initially during the stimulus train, but due to their depression weak facilitating inputs dominate later in the train.

**Figure 2.**
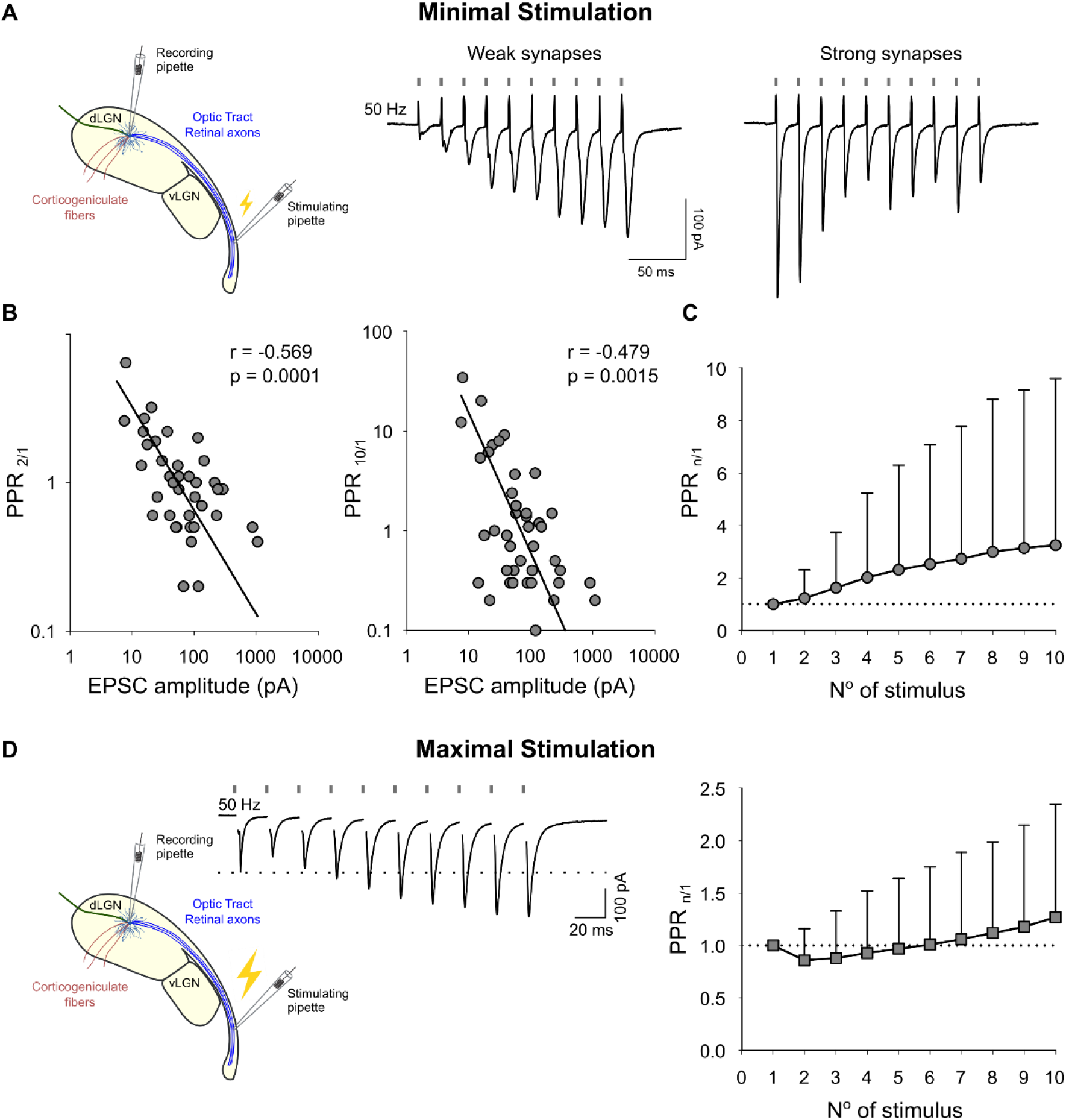
Paired-pulse ratio of retinogeniculate synapses. (A) Schematic representation of minimal stimulation experiments (left). Example currents from strong and weak retinogeniculate synapses. Trains of 10 stimuli delivered at 50 Hz. (B) Dependence of PPR of the second to first response to minimal stimulation on AMPAR-mediated EPSC amplitude (left). (Spearman’s correlation, r=-0.569, ***p=0.0001). The data (black dots) are fitted by a linear regression (black solid line) in log–log space (R^2^=0.537, n=41). Same on the right for the PPR of the tenth to the first response (Spearman’s correlation, r=-0.479, **p=0.0015, R^2^=0.569, n=41). (C) Plot showing the PPR as a function of stimulus number during a 10-pulses train. Each dot represents the mean ± SD across recorded cells. The dotted line marks a ratio of 1 (no facilitation or depression). (D) Schematic representation of maximal stimulation experiments and relative example current (left). On the right, same as in (C), but for maximal stimulation.

### Optogenetic stimulation reproduces short-term plasticity dynamics at retinogeniculate synapses

With electrical stimulation in the optic tract, we targeted axons from RGCs well separated from other inputs to relay neurons, such as from cortex and superior colliculus fibers. Thus, contamination by other inputs with different synaptic properties was unlikely. Nevertheless, to rule out this possibility, we performed control experiments using optogenetic activation of Channelrhodopsin (ChR2)-expressing retinogeniculate axons. For the selective expression of ChR2 in RGCs, we used Chx10-Cre;ChR2 mice, which express Cre recombinase in RGCs but not in neurons from the cortex or superior colliculus^13,25,26^. To achieve stimulation of individual fibers, we restricted the illumination area using a pinhole and gradually increased light intensity from subthreshold levels until minimal, all-or-none responses were detected (Fig. 3A). The distribution of these light-evoked EPSCs was skewed toward small values, closely resembling that of electrically evoked responses (Fig. 3B). On average, the amplitude of light-evoked EPSCs (59.63 ± 96.41; median ± IQR, n=50) was larger than that of electrically-evoked EPSCs. One possible explanation for this would be that it is perhaps easier to activate individual inputs in isolation with electrical stimulation than with light stimulation. Another explanation for the difference in amplitudes is that the vesicle release probability during ChR2-mediated optogenetic stimulation is higher than with electrical stimulation of axons, as shown previously^27^. Indeed, consistent with a higher release probability, PPR of light-evoked EPSCs was lower than that of electrically evoked responses. Importantly, we also observed an inverse correlation between synaptic strength and short-term plasticity in optogenetically evoked responses, both for PPR_2/1_ (Spearman’s correlation, r=-0.384, **p=0.0059) and PPR_10_/_1_ (Spearman’s correlation, r=-0.479, **p=0.0015) (Fig. 3C), as also seen for electrically evoked EPSCs. We then performed maximal stimulation experiments by increasing light intensity and expanding the illumination area to activate multiple fibers simultaneously. This resulted in larger compound EPSCs (402.1 ± 215.0 pA, mean ± SD, n=10). Amplitude and PPR of these compound EPSCs were similar to that of large inputs of minimal stimulation experiments, consistent with a strong contribution of large inputs to compound EPSCs. Together, these findings show that EPSCs evoked by optogenetic stimulation, which potentially increases presynaptic release probability, exhibit strength-dependent short-term plasticity similar to that observed with electrical stimulation.

**Figure 3.**
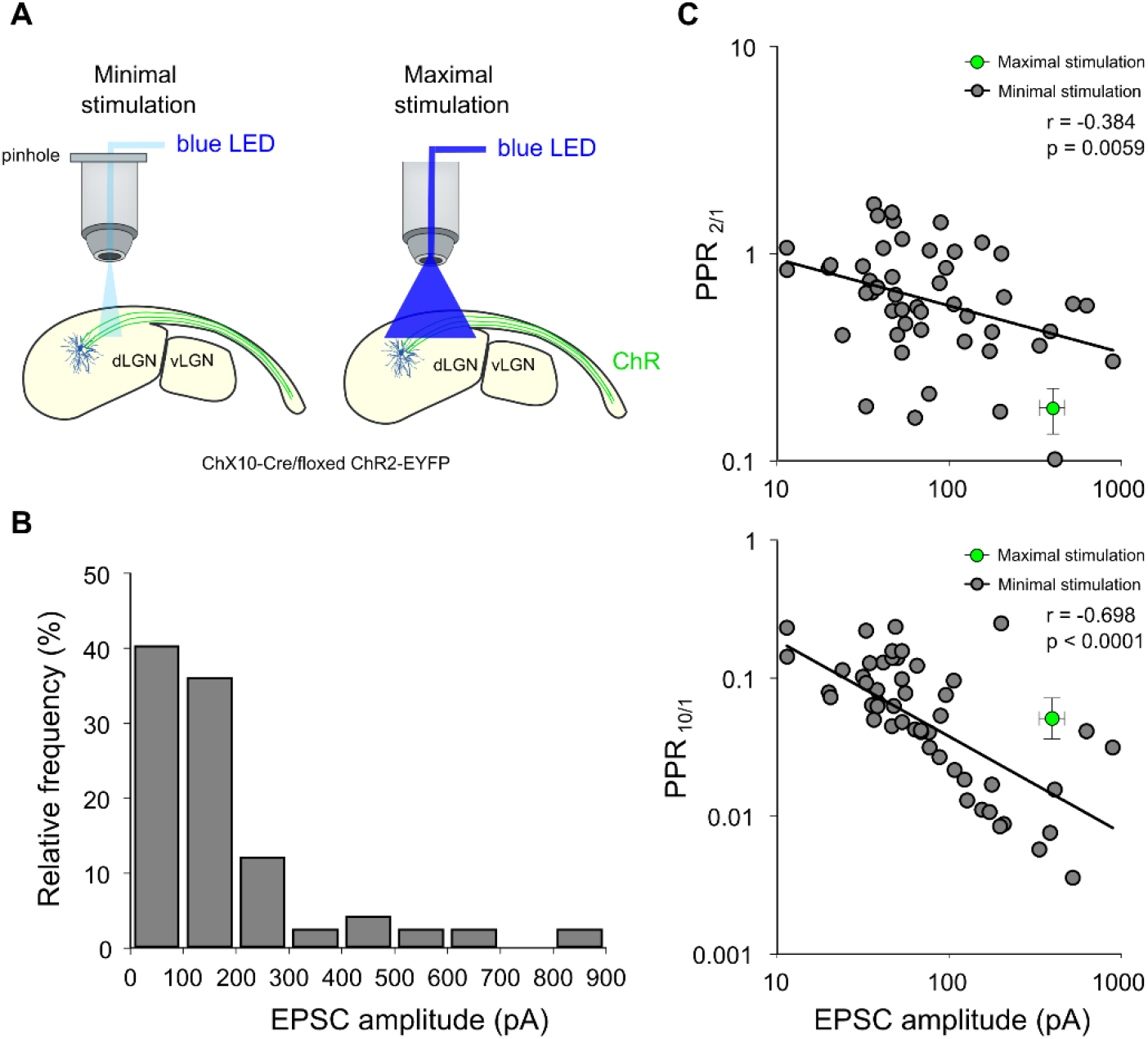
Paired-pulse ratio of retinogeniculate synapses using optical stimulation. (A) Schematic of the experimental setup. Left, for minimal stimulation the LED power was low and the spatial extent of illumination limited via a pinhole. Right, for maximal stimulation the LED power was high and the entire field-of-view was illuminated. (B) Amplitude distribution of first responses to minimal optical stimulation (AMPAR-mediated EPSC amplitude). The distribution is similar to the one obtained with electrically evoked responses, with a strong skew towards small response amplitudes (n=50) (C) 10 stimuli at 50 Hz: dependency of the PPR on the EPSC amplitude. Above, PPR of second to first response to minimal optical stimulation shows a negative correlation (Spearman’s correlation, r=-0.384, **p=0.0059). The data (black dots) are fitted by a linear regression (black solid line) in log–log space (R^2^=0.118, n=50). The green dot is the mean of all responses to maximal stimulation (mean±SD). Below, same but the PPR between the tenth to the first response (Spearman’s correlation, r=-0.698, ****p<0.0001, R^2^=0.309, n=50).

### Presynaptic and postsynaptic mechanisms explain the amplitude dependence of short-term plasticity at retinogeniculate synapses

The inverse correlation between synaptic strength and short-term plasticity might be explained by a lower vesicle release probability in weak than in strong retinogeniculate synapses. Alternatively, short-term depression might increase with synapse strength due to the stronger desensitization of AMPARs in large synapses with many release sites^21^. Finally, both, presynaptic (release probability) and postsynaptic (AMPAR desensitization) mechanisms could explain the inverse correlation. If vesicle release probability increases with synapse strength, one would expect that there is a similar inverse correlation of synaptic strength and short-term plasticity of currents mediated by NMDA receptors, which are not subjected to rapid desensitization. We therefore investigated short-term plasticity of AMPAR- and NMDAR-mediated EPSCs from the same relay cells. EPSCs were evoked by stimulation of the optic tract with 2 pulses at 30 Hz (Fig. 4A). We observed a comparable inverse correlation between synaptic strength and short-term plasticity for AMPAR-mediated (Spearman’s correlation, r=-0.413, **p<0.0001, n=138) and NMDAR-mediated (Spearman’s correlation, r=-0.345, ***p=0.0004, n=103) currents (Fig. 4C), indicating that differences in presynaptic release probability account for a substantial part of this relationship.

**Figure 4.**
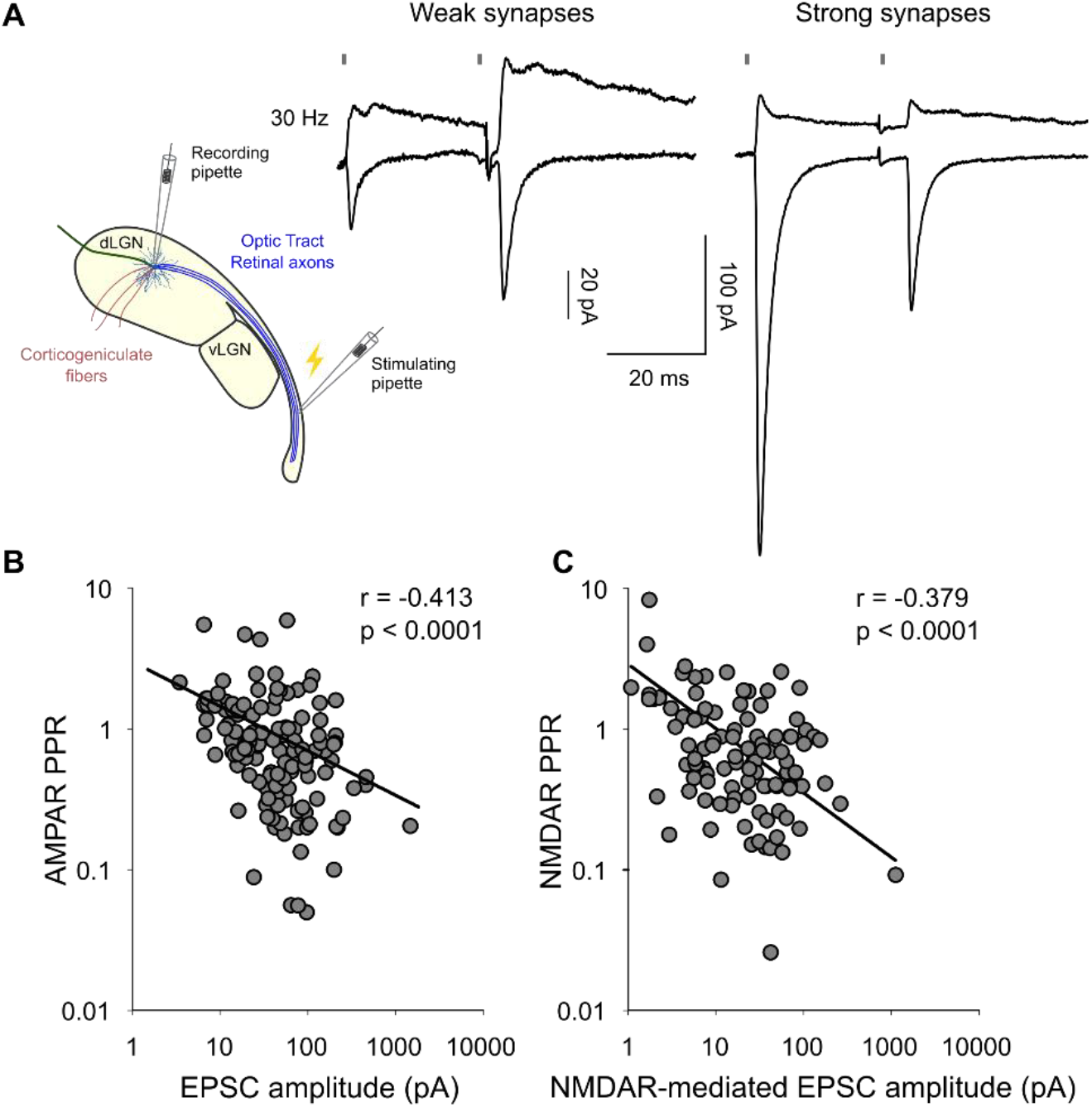
Analysis of AMPAR and NMDAR PPR using minimal stimulation. (A) Schematic representation of minimal stimulation experiments and example currents from strong and weak retinogeniculate synapses. Paired pulses delivered at 30 Hz. (B) Dependence of PPR of AMPAR-mediated current on AMPAR-mediated EPSC amplitude plotted on a logarithmic scale. Negative correlation detected (Spearman’s correlation, r=-0.413, ****p<0.0001). The data (black dots) are fitted by a linear regression (black solid line) in log–log space (R^2^=0.110, n=138). (C) Dependence of PPR of NMDAR-mediated current on NMDAR-mediated EPSC amplitude plotted on a logarithmic scale. Negative correlation detected (Spearman’s correlation, r=-0.345, ***p=0.0004, R^2^=0.226, n=103).

However, these findings do not exclude a contribution of postsynaptic mechanisms. In fact, AMPAR desensitization is expected to be more pronounced in high-release probability than in low-release probability synapses. Moreover, large retinogeniculate synapses with multiple release sites are particularly susceptible to glutamate spillover, which can induce AMPAR desensitization at neighbouring, non-active release sites^21^. To directly assess the contribution of postsynaptic AMPAR desensitization to the inverse correlation of synaptic strength and short-term plasticity, we analysed AMPAR-mediated currents in CKAMP44^‐/‐^ mice. CKAMP44 is an AMPAR auxiliary subunit that enhances receptor desensitization and slows recovery from desensitization^28^. Consistent with our previous work^22^, deletion of CKAMP44 significantly increased PPR during maximal stimulation of retinogeniculate synapses (Mann-Whitney test, n= 19 for wildtype mice, n=22 for CKAMP44^‐/‐^ mice) (Fig. 5A). Importantly, minimal stimulation experiments revealed an inverse correlation between synaptic strength and short-term plasticity in CKAMP44^‐/‐^ mice that was comparable to that observed in wildtype mice (Fig. 5B), supporting the conclusion that presynaptic release probability increases with synapse strength. To determine whether AMPAR desensitization preferentially affects strong synapses, we analysed short-term plasticity separately in weak and strong synapses by using an EPSC amplitude cut-off of 100 pA. PPR was significantly higher in strong retinogeniculate synapses of CKAMP44^‐/‐^ mice compared to the one in wildtype mice (Mann-Whitney test, n=14 for wildtype mice, n=8 for CKAMP44^‐/‐^ mice) (Fig. 5B, right). In contrast, short-term plasticity was not different between genotypes in weak retinogeniculate synapses (Mann-Whitney test, n=27 for wildtype mice, n=26 for CKAMP44^‐/‐^ mice) (Fig. 5B, left). These findings indicate that AMPAR desensitization contributes predominantly to synaptic depression at strong synapses. Together, this suggests that the inverse correlation of synaptic strength and short-term plasticity results from a combination of higher vesicle release probability and more pronounced AMPAR desensitization in stronger compared to weaker retinogeniculate synapses.

**Figure 5.**
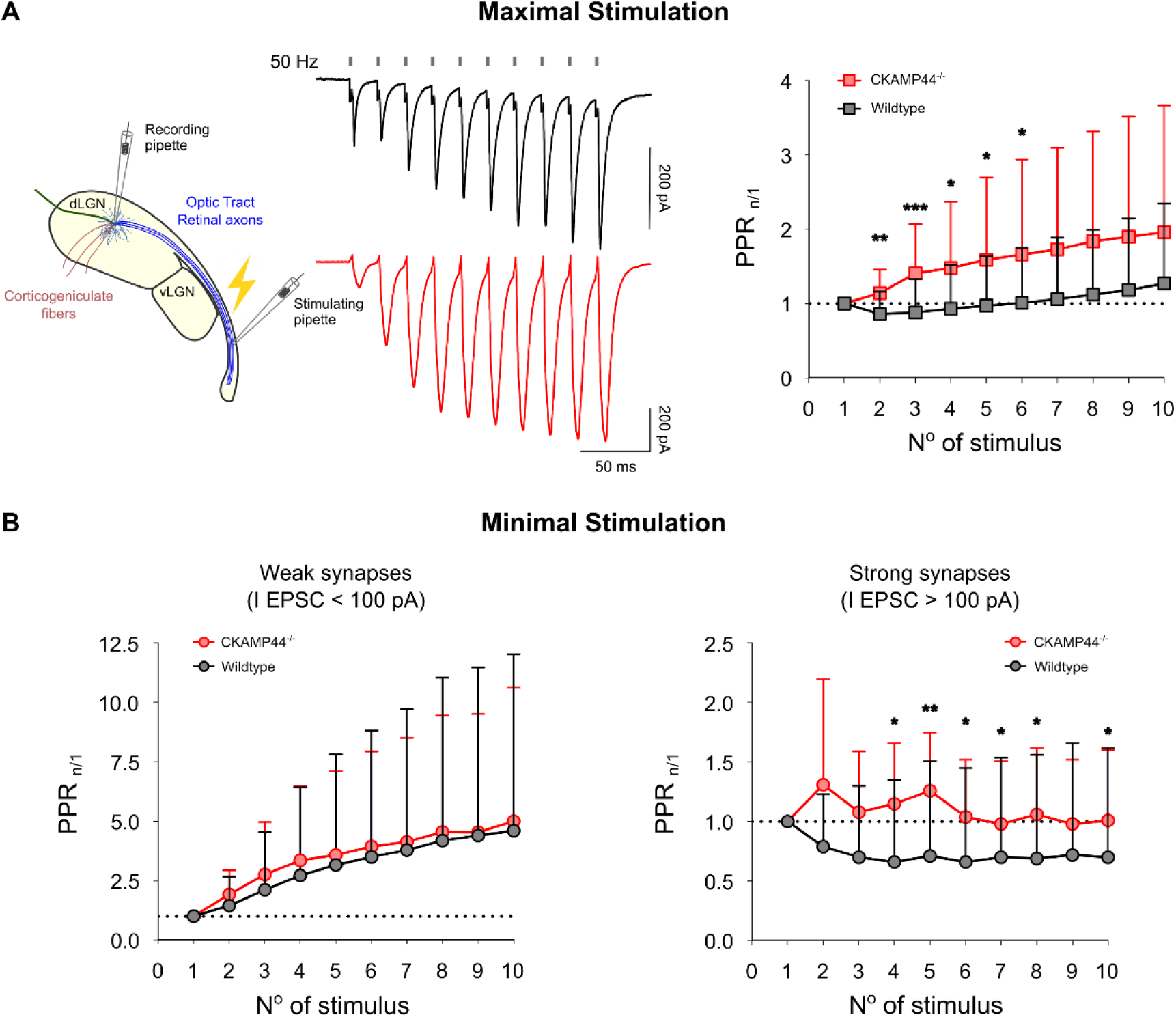
Analysis of PPR of retinogeniculate synapses in wildtype and CKAMP44‐/‐ mice. (A) *Maximal stimulation experiments*. Left, schematic representation of the stimulation setup and representative EPSC traces recorded from wild-type (black) and CKAMP44^-/-^ (red) neurons in response to a 10-pulse train at 50 Hz. Right, PPR plotted as a function of stimulus number during the 10 pulses train. Data are shown as mean ± SD for wildtype (black, n=19) and CKAMP44^-/-^ (red, n=22) neurons. The dotted line marks a ratio of 1 (no facilitation or depression). (B) *Minimal stimulation experiments*. Same as in (A), but retinogeniculate synapses were classified as weak or strong based on the amplitude of the first EPSC (cut-off at 100 pA). Left, weak synapses (first EPSC < 100 pA) showed no significant difference in PPR between genotypes (n=27 for wildtype, n=26 for CKAMP44^‐/‐^). Right, strong synapses (first EPSC > 100 pA) displayed a significant facilitation in CKAMP44^-/-^ neurons and stronger depression in wildtype neurons throughout the stimulus train (n=14 for wildtype, n=8 for CKAMP44^‐/‐^). Data are shown as mean ± SD. *p < 0.05, **p < 0.01, ***p < 0.001

## Discussion

In this study, we provide a detailed analysis of synaptic parameters at retinogeniculate synapses, with a particular focus on the functional differences between weak and strong RGC inputs onto dLGN relay neurons. While the physiological properties of strong retinogeniculate synapses have been extensively investigated^5,13,21,29,30^, considerably less is known about the more numerous weak synapses. Our key finding is that there is an inverse correlation between synapse strength and short-term plasticity, which arises from an interplay between presynaptic and postsynaptic mechanisms. In addition, we found that AMPAR-mediated EPSCs at weak and strong synapses differ in their decay kinetics. This heterogeneity in short-term plasticity and postsynaptic receptor kinetics has important implications for how visual information is transmitted and weighted in the dLGN.

Relay cells in the dLGN receive convergent inputs from multiple RGCs. Functional studies estimate that each relay neuron integrates signals from roughly ten RGCs^13^. In contrast, anatomical studies report substantially higher convergence, identifying up to 91 potential RGCs inputs per relay cell^15,16,31^. However, anatomical approaches cannot assess synaptic strength, and many of the observed contacts may be functionally negligible or even silent (i.e., lacking AMPAR-mediated transmission), rendering them undetectable in physiological recordings. Indeed, the fact that there are many retinogeniculate synapses with EPSC amplitudes just above the noise level suggests that there are also many synapses with EPSCs too small to be detected. During development, most retinogeniculate synapses are comparably weak, with EPSC amplitudes of 10 pA on average^5^. After eye opening, many of those inputs are pruned and the average weight of retinogeniculate synapses increases substantially. However, also in the dLGN of adult mice, the distribution of synaptic strengths is highly heterogeneous with a few strong and many weak inputs^22^.

The EPSC amplitude of single RGC fibers observed in our study was smaller than previously reported^13^. This difference can be well explained by differences in experimental conditions. We used a physiological extracellular Ca^2+^ concentration (1.2 mM) rather than 2 mM Ca^2+^ as described before^13^. When using physiological Ca^2+^, the strength of retinogeniculate synapses is lower than with 2 mM extracellular Ca^2+^ due to a lower vesicle release probability, as shown previously^21^. In addition, we used a K^+^-based intracellular solution instead of cesium. Cesium is known to enhance presynaptic vescicle release probability^32^, and its absence is therefore expected to reduce release probability and increase PPRs, consistently with our previous observations^22^. Accordingly, PPRs of strong inputs obtained from minimal stimulation and PPRs from maximal stimulation experiments in the present study were higher than those reported in studies using 2 mM extracellular Ca^2+^ and a cesium-based intracellular solution^5,13,21,29,30^. Heterogeneity of synaptic strengths could theoretically arise from differences in synapse location along the dendritic tree. Dendritic (electrotonic) filtering reduces the amplitude of EPSCs in distal synapses and slows their kinetics more than those of proximal synapses^24,33,34^. However, the rise time of EPSCs showed a rather weak negative correlation with the amplitude. It is therefore unlikely that different localizations of retinogeniculate synapses contribute substantially to the variability of EPSC amplitudes. However, decay kinetics were negatively correlated with EPSC amplitude. One explanation for this could be that the AMPAR complex composition is different in weak and strong synapses. Indeed, retinogeniculate synapses express several types of GluA subunits, especially GluA1- and GluA4-containing AMPARs^35^. AMPAR homomers exhibit distinct gating kinetics, with GluA4-containing receptors displaying faster deactivation and desensitization compared to GluA1-containing receptors^36^. Differences in the relative contribution of these subunits may influence receptor gating, although AMPAR subunit composition can only explain a small part of the difference in EPSC kinetics. Considering that deactivation kinetics are only marginally affected by the four GluA subunits, auxiliary subunits might play a more prominent role in shaping AMPAR kinetics^37^. Most AMPAR auxiliary subunits, such as TARPs, CNIHs, GSG1l and CKAMPs reduce deactivation kinetics, albeit to different extents^28,38^. Therefore, differential expression and distribution of these auxiliary subunits across retinogeniculate synapses with different weights may contribute to the observed heterogeneity in EPSC kinetics. It could be, for example, that the auxiliary proteins CKAMP44, TARP γ-2 and TARP γ-4, which are all expressed in relay cells and have different effects on current kinetics^22,39^, are differentially expressed in strong and weak synapses.

We observed an inverse correlation of synapse strength and short-term plasticity. The prevailing view is that retinogeniculate synapses display short-term depression. Thus, the facilitation observed in weak synapses was unexpected. However, previous studies did not systematically quantify short-term plasticity of weak inputs using minimal stimulation experiments. When many inputs are activated simultaneously, short-term plasticity is dominated by the depressing strong inputs, masking the facilitation of the weak synapses. Budisantoso and colleagues reported that the short-term plasticity of large inputs is similar to that of weak inputs^21^. However, in this study, individual inputs were not isolated with minimal stimulation, and a high cut-off (500 pA) was used as the criterion for distinguishing between weak and large inputs, effectively comparing large and very large synapses. Under these recording conditions and differentiation criteria for weak and strong synapses, the contribution of truly weak inputs might have been underrepresented.

In control experiments, we confirmed that the inverse correlation between synaptic strength and short-term plasticity is not caused by contamination of non-retinogeniculate inputs (e.g., corticogeniculate synapses). Optogenetic stimulation of retinal axons also showed an inverse correlation of short-term plasticity and synapse strength. Interestingly, PPRs were significantly lower with ChR2 stimulation than with electrical stimulation across the entire spectrum of synapse strengths, suggesting a higher effective release probability during optogenetic activation. This finding is consistent with previous studies showing that ChR2-mediated depolarization enhances presynaptic Ca^2+^ influx and release probability, particularly when stimulation occurs close to presynaptic boutons^27^. Since retinogeniculate axons are bundled in the optic tract, separation of individual fibers was more difficult when stimulating ChR2 far away from the relay cell in the optic tract. Therefore, in our experiments ChR2 stimulation had to be applied near the recorded neuron in the dLGN close to presynaptic terminals, which would further increase release probability and reduce PPR.

The observed negative relationship between AMPAR-mediated synaptic strength and PPR likely arises from strength-dependent differences in both presynaptic and postsynaptic function. Synapse strength increases with the release probability of glutamate vesicles. We found a similar negative relationship for the PPR of NMDAR-mediated currents. Because NMDARs exhibit minimal desensitization during typical transmission, this points to differences in presynaptic release probability as a key contributor to the negative correlation. Our recordings from CKAMP44^-/-^ mice reveal that desensitization also contributes to the negative relationship between synapse strength and AMPAR PPR. CKAMP44 accelerates AMPAR desensitization and slows recovery from desensitization; deletion of this auxiliary subunit increases PPR in retinogeniculate synapses^22^. Here, we show that this effect is limited to strong synapses. The greater involvement of AMPAR desensitization in high-amplitude synapses can be well explained by their higher vesicle release probability^40^. For AMPAR desensitization to influence synapses with a single release site, two consecutive vesicle releases are required. However, retinogeniculate synapses can possess many release sites, allowing glutamate to spillover from active to inactive sites. Consequently, AMPARs desensitize not only at active sites but also at non-active sites due to glutamate spillover^21^. This spillover-dependent desensitization should be more pronounced in synapses with numerous release sites and a high release probability. Therefore, our data suggest that strong synapses exhibit low PPRs because of a combination of high release probability, glutamate spillover, and enhanced AMPAR desensitization.

Although variation in release probability and AMPAR desensitization contributes to overall synaptic strength variability, these factors cannot explain the differences observed among strong synapses (>100 pA), which display similar PPRs and, presumably, comparable release probabilities and desensitization. Instead, the variability in amplitude among these strong synapses is more likely attributable to structural factors such as synapse size, the number of release sites, and the total number of synaptic AMPARs. Consistent with this interpretation, there is no correlation between AMPA/NMDA ratio and synaptic strength. This shows that the number of NMDARs increases with synapse size, similar to the number of AMPARs. Indeed, an electron microscopy study that quantified glutamate receptors in retinogeniculate synapses supports this interpretation, showing that the number of both receptor types increases very similarly with synapse size^41^.

*In vivo* recordings have consistently reported that retinogeniculate synapses exhibit, on average, little short-term plasticity^12,42,43^. This observation has been attributed to a lower release probability *in vivo* compared with *in vitro* conditions, which would reduce the short-term depression at strong synapses. However, our data suggest that this apparent absence of short-term plasticity could emerge from the convergence of a small number of strong depressing inputs and a larger population of weak facilitating inputs onto a single dLGN relay neuron. During high-frequency train activation (50 Hz) of retinal axons, compound EPSCs displayed only a little net change in amplitude. The initial slight depression, which is followed by a slight facilitation can be explained by the dynamic change in the contribution of strong and weak inputs. Strong inputs dominate at the beginning of the EPSC train but rapidly depress, whereas weak inputs progressively increase their contribution. As a result, the total synaptic drive remains relatively stable despite substantial redistribution of synaptic weights across individual afferents. Our data, therefore, indicate that the relative functional contribution of individual RGC inputs is not fixed but depends on the number and frequency of action potentials and therefore on the strength and duration of visual stimuli. This suggests that for short stimuli, firing of relay cells is driven primarily by strong synapses, whereas sustained stimuli recruit weaker facilitating inputs. Since the weak inputs are in the majority, a relay cell may even be triggered to fire primarily by the weak inputs during a long-lasting high-frequency activation. The dynamic adjustment of synaptic strength could aim to stabilize the mean input strength during long-lasting high-frequency activation (activity-dependent gain normalization). This redistribution is expected to have functional consequences for visual coding. If strong and weak inputs convey information from RGCs with similar tuning properties, this dynamic re-weighting would preserve response selectivity. However, anatomical and functional studies demonstrate that a single dLGN neuron receives convergent inputs from multiple RGC types with diverse tuning properties (combination-mode instead of a relay-mode of information transfer)^17,44^. While the input of on average two types of RGCs dominate, the remaining inputs modulate the response of a dLGN neuron^45^. Therefore, changes in the relative synaptic weights of convergent RGC inputs are expected to reshape the tuning properties of relay neurons according to the properties of visual inputs (strength and duration of the visual stimulus) ^14^.

Together, our findings indicate that retinogeniculate synapses are not a homogeneous population and support a model in which they do not simply transmit retinal signals with fixed weights, but might contribute to a dynamic, stimulus-dependent reorganization of information transfer across convergent RGC inputs, providing a flexible structure for adapting visual processing in the dLGN.

## Materials and Methods

### Animals

All animal experiments were performed according to the German Animal Welfare Act and approved by the Governmental Supervisory Panel on Animal Experiments of Rhineland-Palatinate. Mice were bred and raised in-house at the Translational Animal Research Centre (TARC) of the University Medical Centre in Mainz in cages with free access to food and water under a 12-hour light-dark cycle. All animals used in these experiments belong to the C57Bl/6N substrain. CKAMP44^-/-^ mice were generated as described previously^28^. Chx10-Cre;ChR2 mice, which were generated by crossing Chx10-Cre mice (Jax #005105) with Ai32 mice (floxed ChR2-EYFP, Jax #012569), express ChR2-EYFP fusion protein under the control of the Chx10 promoter. Both sexes were used for the experiments.

### Acute Brain Slice Preparation

Acute brain slices containing the optic tract (OT) and dLGN were collected from C57Bl/6N, CKAMP44 ^-/-^or Chx10-Cre;ChR2 (P27 -P50) mice as previously described ^5,22,29^. Briefly, mice were anesthetised with isoflurane and decapitated; the head was cooled, and the brain was removed and immersed in oxygenated ice-cold cutting solution containing (in mM): 110 choline chloride, 3.1 sodium pyruvate, 11.6 sodium-L-ascorbate, 25 NaHCO_3_, 1.25 NaH_2_PO_4_, 25 glucose, 2.5 KCl, 0.5 CaCl_2_, and 7 MgCl_2_. To preserve connectivity between RGC axons and dLGN relay neurons, the two hemispheres were separated with an angle of 3–5° to the sagittal plane and an angle of 18° outwards in the mediolateral plane during slicing. Brain slices (250 μm) were sectioned using a vibratome (Leica VT1200, Leica Microsystems, Wetzlar, Germany), incubated in oxygenated choline-based solution at 34°C for 25 min for recovery and then transferred to artificial cerebrospinal fluid (ACSF) containing (in mM): 125 NaCl, 25 NaHCO_3_, 1.25 NaH_2_PO_4_, 2.5 KCl, 2 CaCl_2_, 1 MgCl_2_ and 25 glucose. Slices were kept at room temperature for about 25 min before electrophysiological recordings.

### In vitro electrophysiology

Whole-cell patch clamp recordings were performed at 34°C using pipettes pulled from borosilicate glass capillaries (Hilgenberg, Malsfeld, Germany) with a resistance of 3–5 MΩ. For recordings of evoked AMPAR-mediated EPSCs, patch pipettes were filled with K^+^-based intracellular solution containing (in mM): 105 K-gluconate, 30 KCl, 10 HEPES, 10 phosphocreatine, 4 Mg-ATP, 0.3 GTP; (pH 7.3, adjusted with KOH). For recording of NMDAR-mediated EPSCs (short-term plasticity and AMPA/NMDA ratios) pipettes were filled with a Cs^+^-based solution containing (in mM): 35 Cs gluconate, 100 CsCl, 10 HEPES, 10 EGTA, and 0.1 D-600; (pH 7.3, adjusted with CsOH). Electrical signals were acquired at 50 kHz using an EPC10 amplifier (HEKA, Reutlingen, Germany) and Patchmaster software (HEKA, Reutlingen, Germany). Liquid junction potentials were not corrected. Series resistance and input resistance were monitored at regular intervals by measuring peak and steady-state current amplitudes in response to small hyperpolarizing voltage steps. Slices were continuously perfused with ACSF containing (in mM): 125 NaCl, 25 NaHCO_3_, 1.25 NaH_2_PO_4_, 2.5 KCl, 1.2 CaCl_2_, 1 MgCl_2_, and 25 glucose; bubbled with 95%O^2^/5%CO^2^ (pH 7.4). Neurons were visualized with an Olympus BX51WI upright microscope (Olympus, Shinjuku, Japan) with a 4x objective (Plan N, NA 0.1; Olympus, Tokyo, Japan) or a 40x water immersion objective (LUMPlan FI/IR, NA 0.8w; Olympus, Shinjuku, Japan) and a CCD camera (XM10, Olympus, Shinjuku, Japan). For electrophysiological patch–clamp experiments, recording and stimulation electrodes were controlled by a SM7 remote unit and control box (Luigs and Neumann, Ratingen, Germany). Recordings of AMPAR-mediated EPSCs were performed in the presence of 50 μM d-2-amino-5-phosphonovaleric acid (d-APV) (Sigma-Aldrich, Taufkirchen, Germany) and 10 μM SR-95531 hydrobromide (gabazine, Biotrend, Cologne, Germany) to block NMDA and γ-aminobutyric acid type A (GABA_A_) receptors, respectively. During recordings of NMDAR-mediated EPSCs, only 10 μM gabazine was added to the ACSF. To evoke EPSCs in retinogeniculate synapses, a monopolar electrode (chlorinated silver wire inside a borosilicate glass capillary filled with ACSF) was placed onto the optic tract. The electrode was connected to a stimulus isolator (A365, World Precision Instruments, Sarasota, FL, USA) that was controlled via Patchmaster software (HEKA, Reutlingen, Germany). Stimulation was repeated 20 times and responses averaged. To identify putative single synaptic connections using minimal stimulation, stimulus intensity was gradually reduced until no EPSCs were detected and then increased until threshold synaptic responses were reliably evoked. Stimulation was then switched from minimal to maximal intensity for each recorded cell and PPR of current amplitudes was calculated from the means of 20 EPSC pairs. To ensure similar stimulation regimes, the stimulation intensity was kept at a similar strength for both genotypes. AMPAR-mediated and NMDAR-mediated EPSCs were recorded at holding potentials of −70 mV and +40 mV, respectively. The optic tract was stimulated with a train of 10 stimuli at 50 Hz for the analysis of PPRs of AMPAR-mediated currents and with two stimuli at 33 Hz for the analysis of PPRs of AMPAR- and NMDAR-mediated currents. PPRs were calculated by dividing the peak amplitude of the n^th^ EPSC by the peak amplitude of the first EPSC. AMPA/NMDA ratios were calculated from the amplitudes of AMPAR-mediated and NMDAR-mediated currents that were recorded at a holding potential of −70 mV and +40 mV, respectively. NMDAR-mediated current amplitudes were measured 25ms after the start of the stimulus artefact. The kinetics of AMPAR-mediated EPSCs were quantified by measuring rise and decay parameters from averaged traces. The 10–90% rise time was defined as the time required for the EPSC to increase from 10% to 90% of its peak amplitude. Decay kinetics were quantified by fitting the falling phase of the EPSC with a double-exponential function of the form: 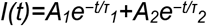, where A_1_ and A_2_ are the amplitudes and τ_1_ and τ_2_ are the fast and slow decay time constants, respectively. A weighted decay time constant (τ_w_) was calculated as: *τ*_*w*_*=(A*_*1*_*τ*_*1*_*+A*_*2*_*τ*_*2*_*)/A*_*1*_*+ A*_*2*_. Fits were performed on the decay phase starting from the peak of the EPSC. Only fits with clear double exponential components and minimal residuals were included in the analysis.

### Optical Stimulation

Optical stimulation experiments were performed on a two-photon microscope (VIVO Multiphoton RS+ with Phasor, 3i). We used a ×16, 0.8 NA microscope objective (Nikon) and the Slidebook Software (3i). Axons were stimulated with a 460 nm LED (pE-300 white, CoolLED) through the objective. For each experiment, axons were stimulated 10 or 20 times. Every trial was 10 seconds long and contained one stimulation. Each stimulation consisted of 10 × 1 ms light pulses at 50 Hz. Stimulation intensity ranged from 1 mW to 100 mW. For minimal stimulation, the light power was defined as the lowest light power that elicited a visible current in the cell. A pinhole was used to further reduce the spatial extent of the stimulation (D25S, Thorlabs). For maximal stimulation the light power was defined as the lowest light power at which the response amplitude was saturated. Signals were synchronized using a data acquisition card (PCI6221, National Instruments) with a BNC break out panel (BNC2090A, National Instruments) and custom-written Labview code (Labview, National Instruments). Patch-clamp recordings were performed as specified above (see *in vitro electrophysiology* section).

### Data Analysis and Statistics

Electrical stimulation experiments were analysed using IGOR Pro 6.37 (WaveMetrics Inc., Lake Oswego, OR, USA) and GraphPad Prism (version 9.3.0, GraphPad Software Inc., San Diego, CA, USA). To assess the relationship between synaptic strength and short-term plasticity, PPRs were plotted against the amplitude of the 1^st^ EPSC in a log-log space. The same approach was applied to AMPAR kinetic parameters. Data were fitted using nonlinear regression with a one-phase decay model. Robust regression was enabled to minimize the influence of outliers and to account for potential non-normality of residuals. Goodness of fit was evaluated using R^2^ values, residual plots, and visual inspection of the fitted curves. Linear correlations were assessed using Pearson’s correlation coefficient for normally distributed data and Spearman’s’s rank correlation coefficient for non-normally distributed data. Statistical comparisons between wildtype and CKAMP44^‐/‐^ mice were performed using an unpaired two-tailed Student’s t-test or Mann–Whitney U-test, depending on data normality, which was assessed using the Shapiro–Wilk test. Data are presented as mean ± standard deviation (SD) for normally distributed datasets or as median with interquartile range (IQR; 25–75%) for non-normally distributed datasets. Sample sizes (n = number of cells) are indicated in the figure legends. Optical stimulation experiments were analysed using custom-written MATLAB scripts. Single traces of each were baseline corrected and averaged. The mean response to all stimulations was smoothed with a gaussian kernel with a span of 0.5 ms, and EPSC amplitudes were extracted as peak values of the mean trace. The minimal stimulation experiments were fitted with an exponential function. P-values <0.05 were considered statistically significant (*p < 0.05, **p < 0.01, ***p <0.001, ****p<0.0001).

## Acknowledgements

We are grateful to B. Biesalski and A. Denner-Seckert for help with animal husbandry and R. Necel for help with equipment. This work was funded by the German Research Foundation (DFG) grant INST 371/52-1, Major Research Instrumentation, 2P-Mikroscope, to J.v.E., and by the DFG grant within the Collaborative Research Centre (SFB) 1080 “Molecular and Cellular Mechanisms of Neural Homoeostasis” to J.v.E. We thank all members and technical staff of the Institute of Pathophysiology for valuable discussions and help.

## Author contributions

J.v.E. acquired funding for the study. F.H., I.S., C.B., and S.R. analyzed the data. J.v.E., C.B., E.J., and S.R., supervised the study. J.v.E., F.H., I.S., E.J., S.R. and C.B. conceived and designed the study. J.v.E., and I.S. wrote the manuscript with critical feedback from all authors.

## Notes

### Competing Interest Statement

The authors have declared no competing interest.

